# Soluble, but not precursor EGF induces intracrine signaling of the EGFR

**DOI:** 10.1101/2025.11.28.691129

**Authors:** Noémi Bilakovics, Julianna Volkó, Chinzorig Jeppesen, Gergely József Fekete, István Csomós, Andrea Bodnár, Gábor Mocsár, Krisztina Németh, Attila Kormos, Katalin Tóth, György Vereb, Péter Nagy, György Vámosi

## Abstract

EGFR is a transmembrane receptor tyrosine kinase regulating growth and survival in epithelial tissues. Its ligand, the epidermal growth factor (EGF), is produced as a membrane-anchored precursor (preEGF) that is proteolytically cleaved to release soluble EGF (sEGF). EGFR overexpression can convert it from a physiological regulator into an oncogenic driver. Therapeutic strategies targeting EGFR include monoclonal antibodies (mAbs) and tyrosine kinase inhibitors (TKIs). Although mAbs such as cetuximab initially block EGFR activity, tumors often develop resistance. Recent findings indicate that sEGF can activate EGFR within intracellular vesicles, promoting intracrine signaling that sustains proliferation despite extracellular inhibition. Here, we investigated the mechanism of intracrine EGFR signaling in the Golgi apparatus using confocal microscopy, FRET and fluorescence correlation spectroscopy. sEGF, but not preEGF, bound and induced EGFR dimerization and phosphorylation. Erlotinib, a membrane permeable TKI, effectively blocked phosphorylation, whereas extracellular cetuximab did not. These findings imply sEGF-induced intracrine EGFR signaling in the Golgi. Our results may shed light on a potential resistance mechanism to antibody treatment.

## Introduction

The Epidermal Growth Factor Receptor (EGFR or ErbB1) is a transmembrane glycoprotein and the first identified member of the ErbB family of receptor tyrosine kinases (Carpenter *et al*, 1978). EGFR plays a pleiotropic role in epithelial tissue cells by inducing multiple intracellular signaling networks responsible for the regulation, development, homeostasis, and proliferation of these cells (Cohen, 1965; Sibilia & Wagner, 1995; Threadgill *et al*, 1995). EGFR comprises three main structural domains: the extracellular domain responsible for ligand binding, the transmembrane domain, and the intracellular domain consisting of the catalytically active tyrosine kinase domain followed by a tyrosine-rich C-terminal tail. (Levkowitz *et al*, 1999; Ohtsu *et al*, 2006; Rotin *et al*, 1992; Sibilia & Wagner, 1995).

EGFR has seven known natural type I transmembrane precursor ligands: epidermal growth factor (EGF), heparin-binding EGF-like growth factor, amphiregulin, epigen, epiregulin, betacellulin and transforming growth factor α. Their structure comprises an N-terminal region, followed by an EGF motif, a juxtamembrane domain, a transmembrane domain and a cytoplasmic tail. Both EGFR and its precursor ligands are synthesized in the ER and follow the same secretory pathway through the Golgi to the plasma membrane. After arriving at the cell membrane, the precursor ligand, such as preEGF, is cleaved releasing soluble sEGF (a process called ectodomain shedding) by the membrane-anchored A Disintegrin And Metalloprotease (ADAM) family enzymes, previously activated via a GPCR-induced process (Daub *et al*, 1996; Ohtsu *et al*., 2006). PreEGF contains nine EGF-motifs in the extracellular region, of which the domain closest to the transmembrane region is shed (sEGF) (Cohen, 1960). Binding of sEGF to EGFR causes dimerization and a conformational change of the receptor chains followed by transphosphorylation of the C-terminal tails (Massague & Pandiella, 1993; Soler *et al*, 1994; Yarden & Schlessinger, 1987). The activation of EGFR leads to the docking of different cytoplasmic regulators and induction of multiple downstream signaling pathways (MAPK, JAK/STAT, PLC-γ and PI3K/Akt) (Anderson *et al*, 1990; Cantley, 2002; Margolis *et al*, 1990; Olayioye *et al*, 1998; Silva, 2004).

EGFR was the first cell surface receptor to be linked to cancer (Downward *et al*, 1984) and it remains an important target in anticancer therapies. Mutations and amplifications in the gene encoding EGFR (Klijn *et al*, 1992; Rubin Grandis *et al*, 1996; Watanabe *et al*, 1996) as well as the presence of autocrine loops (Rubin Grandis *et al*., 1996; Salomon *et al*, 1995; Umekita *et al*, 2000) cause EGFR overexpression in multiple epithelial tumors, such as glioblastoma (Ekstrand *et al*, 1992; Feng *et al*, 2014), breast cancer ( (Nickerson *et al*, 2013; Ro *et al*, 1988; Spyratos *et al*, 1990), non-small cell lung cancer (NSCLC) (Hirsch *et al*, 2003; Hsiao *et al*, 2013; Valley *et al*, 2015) or colorectal cancer (Arena *et al*, 2015).

Over the past decades, scientists have developed inhibitors targeting several key regulatory points in the EGFR signaling pathway to prevent tumor growth, migration, invasion, apoptosis evasion, and metastasis (Cai *et al*, 2020). Two major groups of EGFR inhibitors can be distinguished: extracellularly acting monoclonal antibodies (mAbs) that compete with EGFR ligands for binding, such as cetuximab and panitumumab (Aboud-Pirak *et al*, 1988; Jafary *et al*, 2025; Kimura *et al*, 2007), and cell-permeable small molecule tyrosine kinase inhibitors (TKIs) that block the ATP binding site at the C-terminal region of EGFR and thus the trans-phosphorylation of tyrosine residues, e.g., gefitinib (Iressa) and erlotinib (Tarceva)(Ciardiello, 2000; Cohen *et al*, 2005; Kazandjian *et al*, 2016). The therapeutic action of mAbs involves the promotion of EGFR internalization (Sunada *et al*, 1986) and elimination of tumor cells through antibody-dependent cellular toxicity (ADCC) (Clynes *et al*, 2000; Kimura *et al*., 2007). Despite initial effectiveness, metastatic tumor cells often develop resistance to mAb treatments within months (Neyns *et al*, 2009; Van Cutsem *et al*, 2007; Wheeler *et al*, 2008).

New generation mAbs targeting EGFR are designed to overcome resistance mechanisms and enhance efficacy by inhibiting EGFR more effectively, simultaneously targeting multiple receptors, or through antibody-drug conjugates (ADCs) that inhibit tumor cells both extracellularly via the antibody and intracellularly by releasing the conjugated cytotoxic drug in the cytosol (Jafary *et al*., 2025; Nelhubel *et al*, 2021). Imgatuzumab (GA201), an Fc glycoengineered IgG1 antibody, is optimized to induce both ADCC and EGFR signaling inhibition (Gerdes *et al*, 2013; Kol *et al*, 2017). Another strategy to combat resistance involves (Nelhubel *et al*., 2021) combinations of mAbs and TKIs, such as cetuximab-erlotinib (Huang *et al*, 2004) and amivantamab-lazertinib (Cho *et al*, 2022; Patel & Heppner, 2025), which show promising potential to improve therapeutic outcomes.

Intracrinology, a relatively new field in molecular biology, has identified only a limited number of proteins signaling via this mechanism. The term intracrine (intracellular autocrine) was first introduced in the context of angiotensin II. Angiotensin II is synthesized intracellularly and acts within the nucleus to increase transcription of angiotensinogen and renin (Re & Bryan, 1984). In the case of membrane receptors, the ligand binding ectodomain of a receptor and the ligand are localized in the lumen of the ER, the Golgi and intracellular vesicles along the secretory pathway toward the plasma membrane and may have the opportunity to bind and induce intracrine signaling on route to the cell surface (Brand *et al*, 2011b; Massague & Pandiella, 1993; Volko *et al*, 2019; Yarden & Sliwkowski, 2001). Initially, EGFR ligands were thought to signal exclusively through autocrine, paracrine, juxtracrine, or endocrine pathways (Singh & Harris, 2005). A novel intracrine mechanism was proposed based on observations that soluble EGF colocalized with EGFR within small intracellular vesicles and that phosphorylation could not be blocked by extracellular anti-EGFR antibodies (Wiley *et al*, 1998). However, the molecular details and convincing evidence for the intracrine mechanism are missing.

Several studies have elucidated the molecular mechanisms underlying EGFR function, including an investigation of its diffusion properties (Vamosi *et al*, 2019) and others exploring dimerization/oligomerization and ligand-binding behavior in the presence or absence of external EGF (Clayton *et al*, 2007; Clayton *et al*, 2005; Hajdu *et al*, 2020; Lidke *et al*, 2004; Needham *et al*, 2016; Zanetti-Domingues *et al*, 2018). In the past years, our research group focused on the intracrine signaling of IL-2-receptor discovering that the IL-2 receptor heterotrimer preassembles in the ER and the Golgi before reaching the cell membrane and phosphorylation of signaling components occurs already in the Golgi of adult T lymphoma cells co-expressing IL-2 and IL-2R (Volko *et al*., 2019). These cells also evaded blocking by antiproliferative antibodies. We were interested in finding out whether the steps of this intracellular signaling mechanism could be observed and defined for the EGFR in the Golgi by using fluorescence microscopy techniques. Chinese hamster ovary (CHO) cells lack endogenous EGFR (Shi *et al*, 2000) making it an ideal model system for expressing our proteins of interest. We investigated sEGF/preEGF ligand binding and receptor chain dimerization in the Golgi apparatus using fluorescence lifetime imaging microscopy-based Förster resonance energy transfer (FLIM-FRET) and fluorescence correlation spectroscopy (FCS). EGFR activation was detected by an anti-pEGFR mAb specific for the phosphorylated Y1173 residue of the C-terminal tail. We also assessed the inhibitory effects of the cetuximab mAb and the intracellular TKI erlotinib on EGFR phosphorylation in externally EGF-treated as well as in sEGF-producing cells. We showed that if the tumor cell expresses both EGFR and sEGF ligand, intracrine signaling takes place. Our results suggest that one of the underlying reasons for resistance to mAb therapies in tumors overexpressing EGFR and producing EGF simultaneously may be intracrine EGFR signaling.

## RESULTS

### FLIM-FRET indicates sEGF ligand binding to EGFR in the Golgi

FLIM-FRET is an ideal tool for measuring molecular protein-protein interactions such as receptor-ligand binding or receptor dimerization. In this study, we performed FLIM-FRET imaging to examine the binding of soluble and precursor EGF (sEGF, preEGF) ligands to EGFR in the Golgi apparatus. The donor was tagged with eGFP (abbreviated G), while the acceptor was mScarlet3 (S). FRET occurs when the donor is within 2–10 nm proximity of the acceptor, leading to a shortening in the donor’s fluorescence lifetime, which forms the basis of FRET efficiency (E) calculations (*see* *equation 8*).

Confocal images of selected cells were captured in the donor (G), acceptor (S), Golgi marker (B-Giantin) channels as well as the transmission (TM) channel (*Fig. 1A*, *columns 1–3*) prior to FLIM measurements. Merged images of the donor, acceptor and Golgi marker channels are also shown (*Fig. 1A*, *column 4*). FRET efficiencies were assessed either across the whole cell or specifically in the pixels positive for the B-Giantin Golgi marker. Corresponding FRET efficiency maps and histograms are shown on *Fig. 1A, columns 5-6*. The average E% for each sample is shown on a scatter dot plot on *Fig. 1B*. Schematic drawings of the measured control samples (eGFP-mScarlet3 fusion (abbreviated G-S), G and S, G-sEGF and S-IL-2Rα (interleukin-2 receptor α chain)), the G-sEGF and S-EGFR, S-preEGF and G-EGFR or preEGF-S and EGFR-G are shown on *Fig. 1C*.

**Fig. 1.**
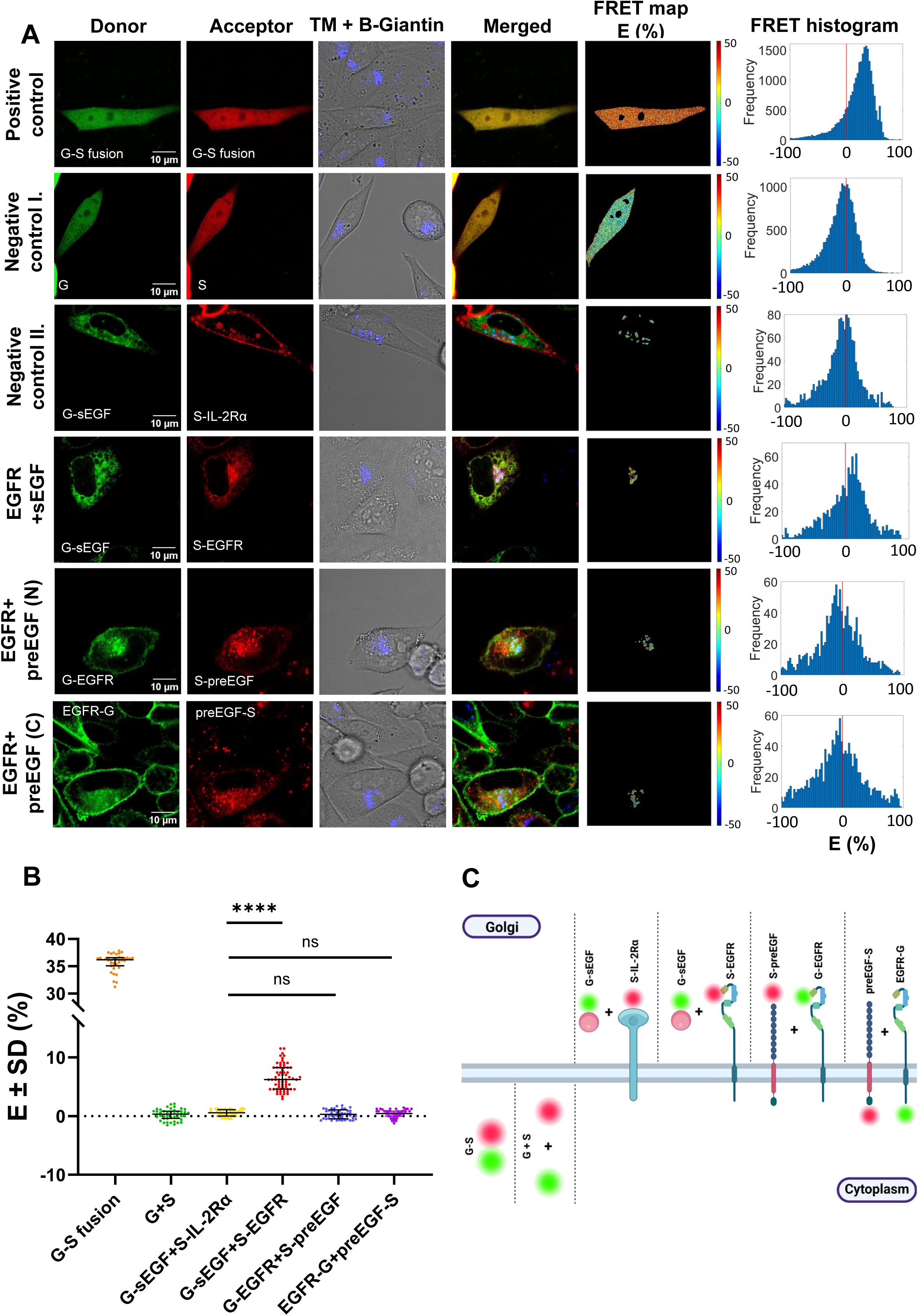
FLIM-FRET measurement of the assembly of EGFR and preEGF/sEGF proteins in the Golgi-apparatus in CHO and F1.4 cells. **(A)** Confocal images of the FRET donor (1^st^ column), acceptor (2^nd^ column), transmission image with the BFP labelled Golgi (3^rd^ column), merge of all channels (4^th^ column) and FRET efficiency maps/histograms (5^th^-6^th^ columns) of selected cells. On the histograms the x axis shows the FRET efficiencies in %, the y axis shows the frequency values. The fluorescent protein tags were abbreviated as B: TagBFP, G: eGFP, S: mScarlet3). 1^st^ row: positive control, G-S fusion protein; 2^nd^ row: negative control I, co-transfected G and S constructs; 3^rd^ row: negative control II, G-sEGF co-transfected with S-IL-2Rα; 4^th^ row: G-sEGF donor expressed with S-EGFR; 5^th^ row: N-terminally G- and S-tagged EGFR and pre-EGF, respectively; 6^th^ row: C-terminally G- and S-tagged EGFR and preEGF. The scale bar is 10 µm. **(B)** The graph shows the average FRET efficiencies and standard deviations of the samples; the significant increase in *E* (relative to the negative control II.) of G-sEGF with S-EGFR indicates their association in the Golgi. Unpaired t-tests were performed to compare averages. ns: not significant; ****: p<0.0001. N=38-67 cells were measured for each sample. **(C)** Schematic drawing of the measured samples: positive control (G-S fusion), negative control I. (G and S), negative control II. (G-sEGF+ S-IL-2Rα); G-sEGF and S-EGFR; S-preEGF and G-EGFR; preEGF-S and EGFR-G.

We used the homogeneously distributed G-S fusion protein as a positive control, which had an average FRET efficiency of E = 35.7 ± 1.6% (mean ± SD) (*Fig. 1A, first row; Fig. 1B*). With the help of this fusion protein the donor-to-acceptor molecular ratios could be determined for each donor-acceptor double-transfected sample; we selected cells around or above a 1:1 donor/acceptor ratio. Next, we assessed two different negative controls. First, for negative control I, G and S were transiently co-transfected and distributed evenly in the whole cell yielding E=0.2±0.9% (*Fig. 1A, 2^nd^ row; Fig. 1B*). Second, G-sEGF and an irrelevant receptor, S-IL-2Rα were co-expressed resulting in E=0.6±0.5% in the Golgi (*Fig. 1A, 3^rd^ row; Fig. 1B*). This second negative control demonstrates the level of random FRET that may occur between two non-related proteins in the Golgi apparatus. We selected this control as a baseline to compare FRET efficiencies of the sEGF/preEGF and EGFR-expressing cells. Co-expression of G-sEGF and S-EGFR resulted in E=6.5±2.1% in Golgi marker positive pixels (*Fig. 1A, 4^th^ row; Fig. 1B.*), which is significantly higher than the FRET efficiency in negative control II. This elevated E level indicates sEGF binding to EGFR in the Golgi. Next, G-EGFR was transfected into CHO cells stably expressing S-preEGF and B-Giantin, yielding E=0.2±0.8% (*Fig. 1A, 5^th^ row; Fig. 1B*). The measurement was also performed with the C terminally tagged preEGF-G and EGFR-S, where the F1.4 cells expressing EGFR-G were stably transduced with preEGF-S and B-Giantin; the FRET efficiency was E=0.5±0.7% for this sample (*Fig. 1A, 6^th^ row; Fig. 1B*). Since neither of these FRET efficiency values differed significantly from that of negative control II, we can assume that there is no significant association between preEGF and EGFR in the Golgi.

The FRET efficiency was plotted as a function of the acceptor-to-donor molecular ratio (N_a_/N_d_) calculated according to *Equations 9 and 10* for all samples (*Fig. S1*). In the sEGF and EGFR expressing sample E showed an increasing tendency with an increasing N_a_/N_d_ ratio (Fig. S1C). As the N_a_/N_d_ ratio increases, donors are more likely to pair with acceptors enabling FRET and increasing the average FRET efficiency as expected. In contrast, all the other samples exhibited no consistent tendency (*Fig. S1A, B, D* and *E*). The fluorescence lifetime decay curves of the selected cells shown in *Fig. 1A*, fitted to a two-component Exponential Reconvolution model are shown in *Fig. S2A*.

### The presence of EGFR causes a decrease in sEGF mobility within the Golgi

We investigated ligand binding in the Golgi by using FCS to measure the mobility of G-sEGF in the presence or absence of S-EGFR. Two control samples were included in the measurement: free eGFP and membrane-bound eGFP-EGFR. Constructs were transiently transfected into live CHO cells stably expressing the B-Giantin Golgi marker 24 h prior to the measurement. To ensure equal expression of the fluorescent proteins, the amounts of transfected DNA were optimized, and the resulting intensity ratios were compared with those of a G–S fusion protein, which provides a defined 1:1 expression ratio. *Fig. 2A* shows representative confocal images of selected cells, where B-Giantin, G-sEGF and S-EGFR or their merged view can be seen.

**Fig. 2.**
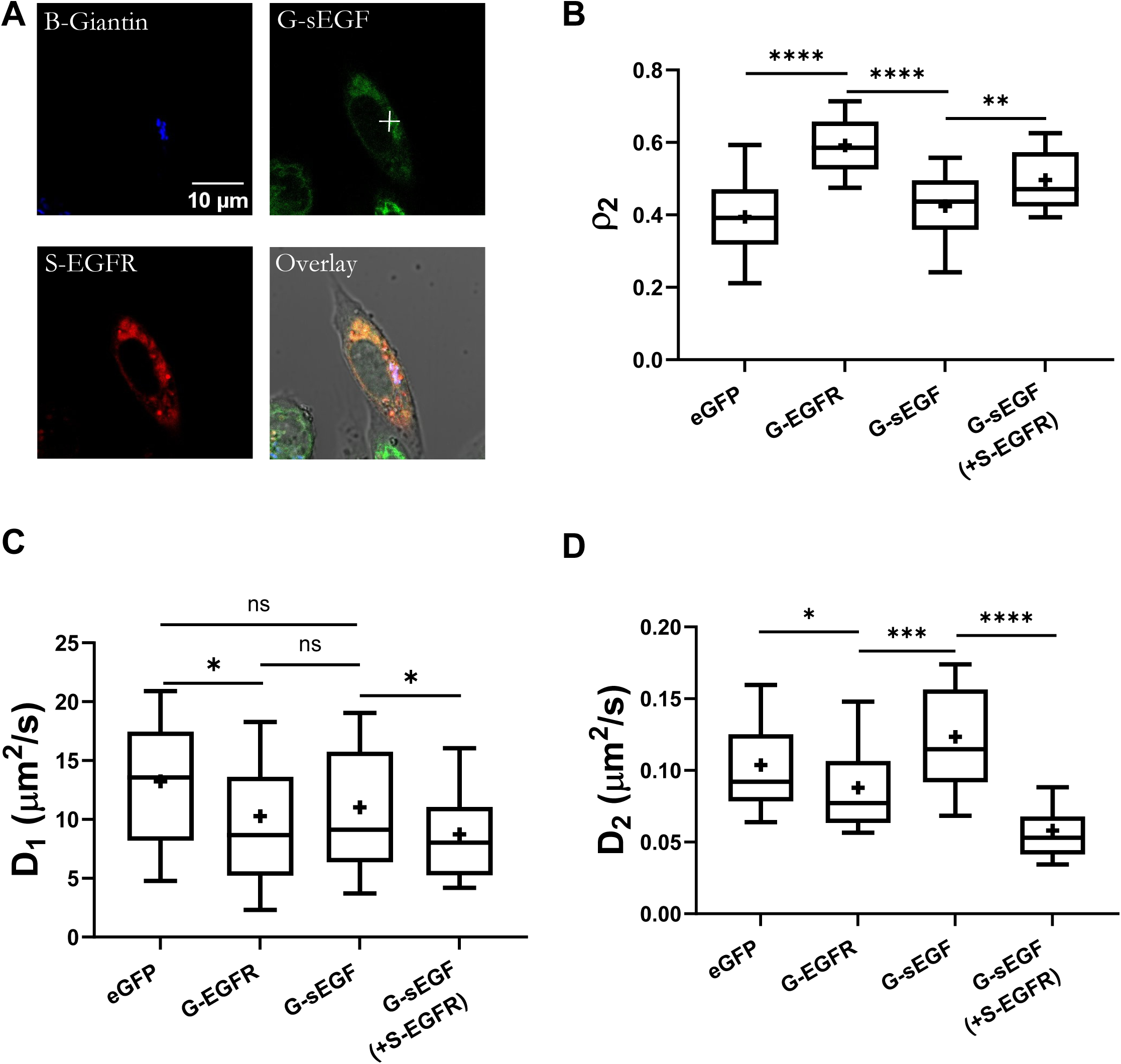
Representative confocal image and mobility parameters of eGFP, G-EGFR, G-sEGF expressed alone and G-sEGF coexpressed with S-EGFR in the Golgi of CHO cells. **(A)** Confocal images of the B-Giantin, G-sEGF and S-EGFR in live CHO cells. The composite image of all channels shows the overlay of G-sEGF and S-EGFR in the Golgi with white color. The white cross in the G-sEGF channel indicates the point where the FCS measurement took place. The scale bar is 10 µm. **(B)** The fraction of the slow component (ρ2) of eGFP, G-EGFR, G-sEGF proteins alone and G-sEGF co-transfected with S-EGFR. **(C)** Diffusion coefficients of the fast component (D1) of eGFP, G-EGFR, G-sEGF (alone or in the presence of S-EGFR). **(D)** Diffusion coefficients of the slow component (D2) of eGFP, G-EGFR and G-sEGF (alone or in the presence of S-EGFR). The eGFP, G-sEGF and G-sEGF/S-EGFR were fitted with a two-component 3D normal model, while G-EGFR with a 2D normal model. The horizontal line in the middle represents median values, and the bottom and top of the box show the 25th and 75th percentiles, while the whiskers indicate the 10th and 90th percentiles. Averages are shown with a ‘+’ sign in the boxes. Unpaired t-test was performed to compare sample averages. ns: not significant, *: p<0.05,**: p <0.01, ***: p<0.001, ****: p<0.0001 (t-test). N=34-53 cells were measured for each sample.

In FCS, the fluctuating fluorescence intensity of the eGFP tag at a selected point of the Golgi was measured and its autocorrelation function (ACF) was calculated. ACFs were fitted with two-component 2D or 3D normal diffusion models (*Fig. S3*). A 2D model was used only for G-EGFR because it is a membrane-bound protein, for which diffusion is restricted to 2D; the diffusion of the ligand was fitted to a 3D model. The fit yields the fractions of the fast and slow components (ρ₁ and ρ₂) and the diffusion coefficients (D₁ and D₂, in µm²/s) or the diffusion (dwell) times (τ_D1_ and τ_D2_, in µs). The diffusion coefficient is inversely proportional with the diffusion time (*See equations 2 and 3*). Here, we only show the D_1_, D_2_ and ρ_2_ parameters (*Fig. 2B, 2C, 2D*).

First, we measured the mobility parameters of eGFP in the Golgi, which serves as a freely diffusing inert, non-binding control protein (Vamosi *et al*, 2016). The diffusion properties of G-EGFR were also investigated previously (Vamosi *et al*., 2019) but only in the cell membrane, not the Golgi. The ρ_2_ slow fraction was 0.39±0.13 (mean±SD) for free eGFP, while it was significantly higher, 0.59±0.08 for G-EGFR in the Golgi (*Fig. 2B*). This reflects the more restricted mobility of EGFR due to its membrane-bound nature compared to more freely diffusing eGFP. For G-sEGF expressed alone, ρ_2_ was 0.42±0.12, which is significantly lower than for G-EGFR indicating that sEGF also diffuses more freely in the Golgi than EGFR. The fraction of the slow component of G-sEGF increased to 0.49 ± 0.09 when S-EGFR was present, indicating that EGFR binds sEGF.

The average D_1_ diffusion coefficient of the fast component was significantly larger for eGFP (D_1_=13.2±5.97 µm^2^/s) than for G-EGFR (10.3±8.1 µm^2^/s), while for G-sEGF (11.0±5.9 µm^2^/s) not significant (*Fig. 2C*). The D_1_ value of G-sEGF significantly decreased when S-EGFR was co-expressed (8.8±4.1 µm^2^/s). Comparison of the slow diffusion coefficients (D_2_) shows that the membrane resident G-EGFR has a significantly lower mobility (D_2_=0.08±0.03 µm^2^/s) than free eGFP (D_2_=0.10±0.04 µm^2^/s) or G-sEGF expressed alone (D_2_=0.12±0.04 µm^2^/s). When S-EGFR is co-expressed with G-sEGF, the mobility of the ligand significantly decreases (to 0.058±0.02 µm^2^/s) (*Fig. 2D*). Taken together, all three diffusion parameters imply that sEGF diffuses in the Golgi more slowly when EGFR is present, which suggests that at least a fraction of the ligands is in a receptor-bound state.

### EGFR dimerization in the Golgi is induced by sEGF, but not by preEGF

To assess the association between EGFR-G (donor) and EGFR-S (acceptor) without or with preEGF/sEGF ligand we used FLIM-FRET. We used C-terminally tagged versions of the EGFR chains because the intracellular C-terminal domain is structurally less complex than the extracellular region, which may allow the attached fluorescent dyes to come into closer proximity. To visualize the Golgi, Giantin carrying an N-terminal HaloTag was expressed and labelled with a near-infrared SiR HaloTag ligand (Lukinavicius *et al*, 2013). Representative confocal images of the BFP-tagged ligands preEGF and sEGF (Channel 1), the EGFR-G donor (Channel 2), the EGFR-S acceptor (Channel 3), SIR-labelled HaloTag-Giantin (H-Giantin) together with the transmitted light (TM) image (Channel 4) and the merge of the fluorescence channels (1, 2 and 4) are shown (*Fig. 3A, columns 1-5*). In the merged image the magenta color of H-Giantin was changed to red to highlight the triple overlap by white color. FRET efficiencies were evaluated for the entire cell in case of the G-S fusion. In the absence of ligand or in the presence of preEGF, FRET was assessed both in the Golgi and in the cell membrane. For sEGF, the receptor was located in intracellular vesicles/Golgi, just a few cells expressed the EGFR partially in the plasma membrane; therefore, the evaluation was only applied in the Golgi. In cells treated with external EGF (extEGF), FRET was analyzed only in the cell membrane. FRET efficiency maps and histograms (E (%)) of the selected cells are shown in *Fig. 3A, columns 6-9.* The fitted fluorescence lifetime decay curves of the selected cells in *Fig. 3A* are shown in *Fig. S4*.

**Fig. 3.**
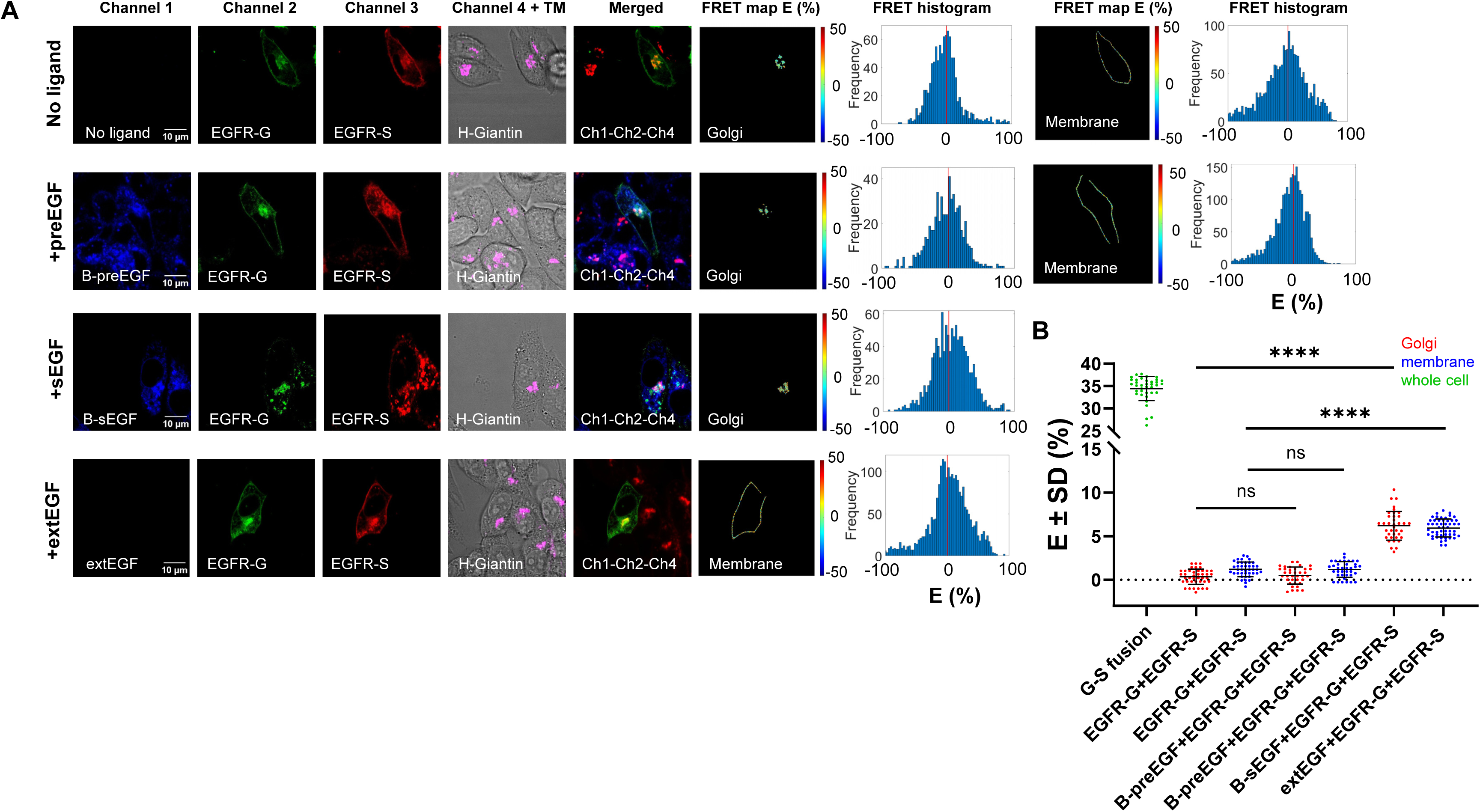
FLIM-FRET measurement of EGFR dimerization in the Golgi-apparatus or the membrane of live CHO cells. **(A)** Representative confocal images of non-transfected, B-preEGF or B-sEGF transfected and externally EGF treated (extEGF, 200 nM) CHO cells in channel 1 (1^st^ column), the FRET donor (EGFR-G) in channel 2 (2^nd^ column), the acceptor (EGFR-S) in channel 3 (3^rd^ column), Halotag-Giantin (H-Giantin) Golgi marker with the TM image (4^th^ column), merge of Ch1-Ch2-Ch4 channels (5^th^ column); FRET effiency map and histogram in the Golgi or in the cell membrane (6^th^-9^th^ columns). On the histograms the x axis shows the FRET efficiencies in %, the y axis shows the pixel frequencies. In the merged image the magenta pseudocolor of H-Giantin was changed to red to see the overlap in white. 1^st^ row: EGFR-G+EGFR-S (no ligand); 2^nd^ row: B-preEGF+EGFR-G+EGFR-S; 3^rd^ row: B-sEGF+EGFR-G+EGFR-S; 4^th^ row: extEGF (200 nM)+EGFR-G+EGFR-S. The scale bar is 10 µm. **(B)** Average FRET efficiencies (mean±SD (%)) of the samples. G-S fusion was measured in the whole cell, dimerization was measured on the Golgi (marked red) or in the plasma membrane (blue). The horizontal lines represent the averages and the standard deviations. Unpaired t-tests were performed to compare averages; ns: not significant; ****: p<0.0001. N= 37-52 cells were measured per sample.

Average FRET efficiencies in different cell compartments are shown in *Fig. 3B*. 37-52 cells were measured for each sample. G-S fusion was used as a positive control, where an average FRET efficiency of E= 34.5± 2.7% (mean ± SD) was measured. EGFR-G co-expressed with EGFR-S without any ligand served as a baseline for subsequent experiments in the presence of ligands (*Fig. 3A, 1^st^ row*). For this sample, in the Golgi, E=0.4±0.9%, while in the plasma membrane, E=1.2±0.8% was detected (*Fig. 3B*). The same tendency was observed in cells stably transfected with B-preEGF (*Fig. 3A, 2^nd^ row*): E=0.5±1.0% was measured in the Golgi, and a slightly higher, E= 1.2±0.9% value was detected in the plasma membrane (*Fig. 3B*). These values did not differ significantly from those in the corresponding no-ligand samples. The slightly higher FRET efficiency in the cell membrane might imply the presence of pre-formed EGFR dimers independently from any ligand, as was reported previously (Hajdu *et al*., 2020). When B-sEGF was co-expressed with EGFR-G and EGFR-S, the FRET efficiency in the Golgi increased to E=6.2±1.7%, which is significantly higher than in non-treated or in preEGF-expressing cells (*Fig. 3A, 3^rd^ row; Fig. 3B*). These results suggest that at least partial homoassociation of EGFR chains occurs in the presence of sEGF in the Golgi. As a positive control, cells co-expressing EGFR-G and EGFR-S were treated with 200 nM extEGF ligand for 5 minutes, resulting in a FRET efficiency of E=5.9±1.0% in the plasma membrane, which is significantly higher than in untreated cells, indicating ligand-induced receptor activation (*Fig. 3A*, *4^th^ row; Fig. 3B*).

The FRET efficiency was plotted as a function of the acceptor-to-donor ratio (N_a_/N_d_) for all samples (*Fig. S5*). The N_a_/N_d_ ratios in the samples were calculated according to *Equations 9 and 10*. As expected, *E* increased in the Golgi with increasing N_a_/N_d_ ratio in the presence of sEGF, and higher FRET efficiency was observed in the high donor expression group than in the low donor expression group (*Fig. S5E*). The same tendency was observed in the extEGF-treated sample (*Fig. S5F*). No tendency was observed in FRET efficiency changes for the non-treated or preEGF expressing cells in the Golgi (*Fig S5A, C)*.

### sEGF activates EGFR in the Golgi, while preEGF does not

To test whether preEGF/sEGF induces receptor phosphorylation, we used immunofluorescent staining of pEGFR in CHO cells and F1.4 cells expressing B-Giantin. The monoclonal antibody recognizes the phosphorylated Y1173 residue of EGFR. Representative confocal images show the distribution of B-Giantin (Ch1), EGFR-G (Ch2), the S-preEGF or S-sEGF ligands (Ch3), the anti-pEGFR-mAb conjugated with AlexaFluor-647 (Ch4) and the merge between channels 1, 3 and 4 (*Fig. 4A*). EGFR was localized in the plasma membrane in the preEGF expressing sample, and it was similar to control F1.4-BFP-Giantin cells. The anti-pEGFR mAb gave a homogenous, low-intensity background signal only. In cells transfected with sEGF, the distribution of EGFR changed completely, it was localized dominantly in intracellular vesicles including the Golgi. The phosphorylation level of EGFR was high in these cells, indicating ligand binding and receptor activation in the Golgi (*Fig. 4A*).

**Fig. 4.**
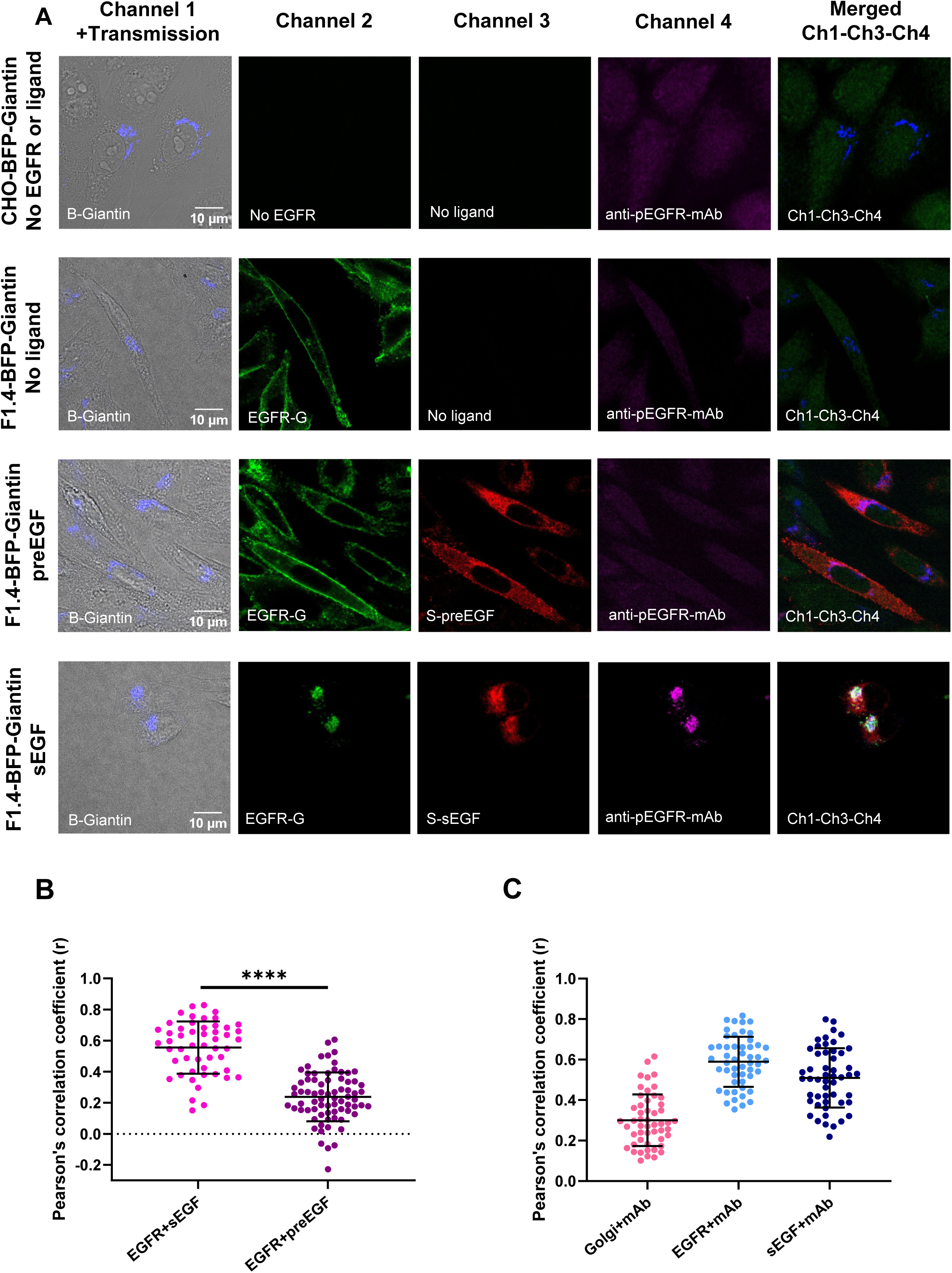
Immunofluorescence labelling of pEGFR shows activation by sEGF but not by preEGF in F1.4-BFP-Giantin cells. **(A)** Confocal images of selected cells. 1^st^ column: the B-Giantin Golgi marker overlayed with the transmission channel, 2^nd^ column: EGFR-G (absent in the top row); 3^rd^ column: transfected preEGF-S (3^rd^ row) or sEGF-S (4^th^ row) ligands (absent in the 1^st^ and 2^nd^ rows), 4^th^ column: Alexa 647-tagged anti-pEGFR-mAb; 5^th^ column: overlay of Ch1-Ch3-Ch4 (Giantin, ligand, pEGFR) without the transmission image. In the merged image the pseudocolor of the anti-pEGFR-mAb channel was changed from magenta to green to see the triple overlap in white color. 1^st^ row: control CHO cell line; F1.4 cells stably expressing B-Giantin with no ligand present (2^nd^ row), expressing preEGF (3^rd^ row) or sEGF (4^th^ row). The scale bar is 10 µm. **(B)** Pearson’s correlation coefficients were calculated between EGFR + sEGF or EGFR + preEGF pairs in the Golgi. **(C)** Pearson’s correlation analysis of Golgi+anti-pEGFR-mAb, EGFR+pEGFR-mAb and sEGF+anti-pEGFR-mAb pairs in F1.4-BFP-Giantin cells. The horizontal lines represent the averages and the standard deviations of the samples. To compare the averages unpaired t-test was used; ****: p<0.0001. N=51-74 cells were measured per sample.

We performed cell-by-cell Pearson’s correlation analysis to quantify the colocalization between different markers within the area of the Golgi as defined by B-Giantin positive pixels. Correlation coefficients were assessed between EGFR and the ligands resulting in r=0.56±0.16 (mean±SD) for EGFR+sEGF and 0.24±0.15 for EGFR+preEGF (*Fig. 4B*). The significantly higher correlation coefficient for sEGF suggests that it is colocalized with the receptor in the Golgi, contrary to preEGF. We also calculated the correlation coefficients for the codistributions of the anti-pEGFR mAb with B-Giantin, EGFR-G or S-sEGF in sEGF-producing cells; in the preEGF-expressing cells this was not possible because of the very low pEGFR signal. For B-Giantin+anti-pEGFR-mAb pair we got r=0.3±0.13, for EGFR+anti-pEGFR-mAb, r=0.59±0.12 and for sEGF+anti-pEGFR-mAb, r=0.51±0.14, indicating a high degree of colocalization of these protein pairs (*Fig. 4C*). Taken together, our data imply that sEGF binds to EGFR in the Golgi and induces receptor phosphorylation, while preEGF does not cause activation.

### Effect of intracellular and extracellular inhibitors on EGFR phosphorylation

Two inhibitors, the extracellularly blocking cetuximab and intracellularly acting kinase inhibitor erlotinib were tested for their effect on EGFR phosphorylation. Two samples were used: F1.4 cells pretreated with inhibitors and then treated with 200 nM extEGF for 10 minutes or F1.4 cells transfected with S-sEGF and treated with inhibitors (*Fig. 5*). After the treatments, cells were labelled with AlexaFluor-647-tagged anti-pEGFR-mAb and confocal images were recorded in the EGFR-G, S-sEGF, and anti-pEGFR-mAb channels. Representative images of cells with different treatments are shown in *Fig. S6A* and *S7A*.

**Fig. 5.**
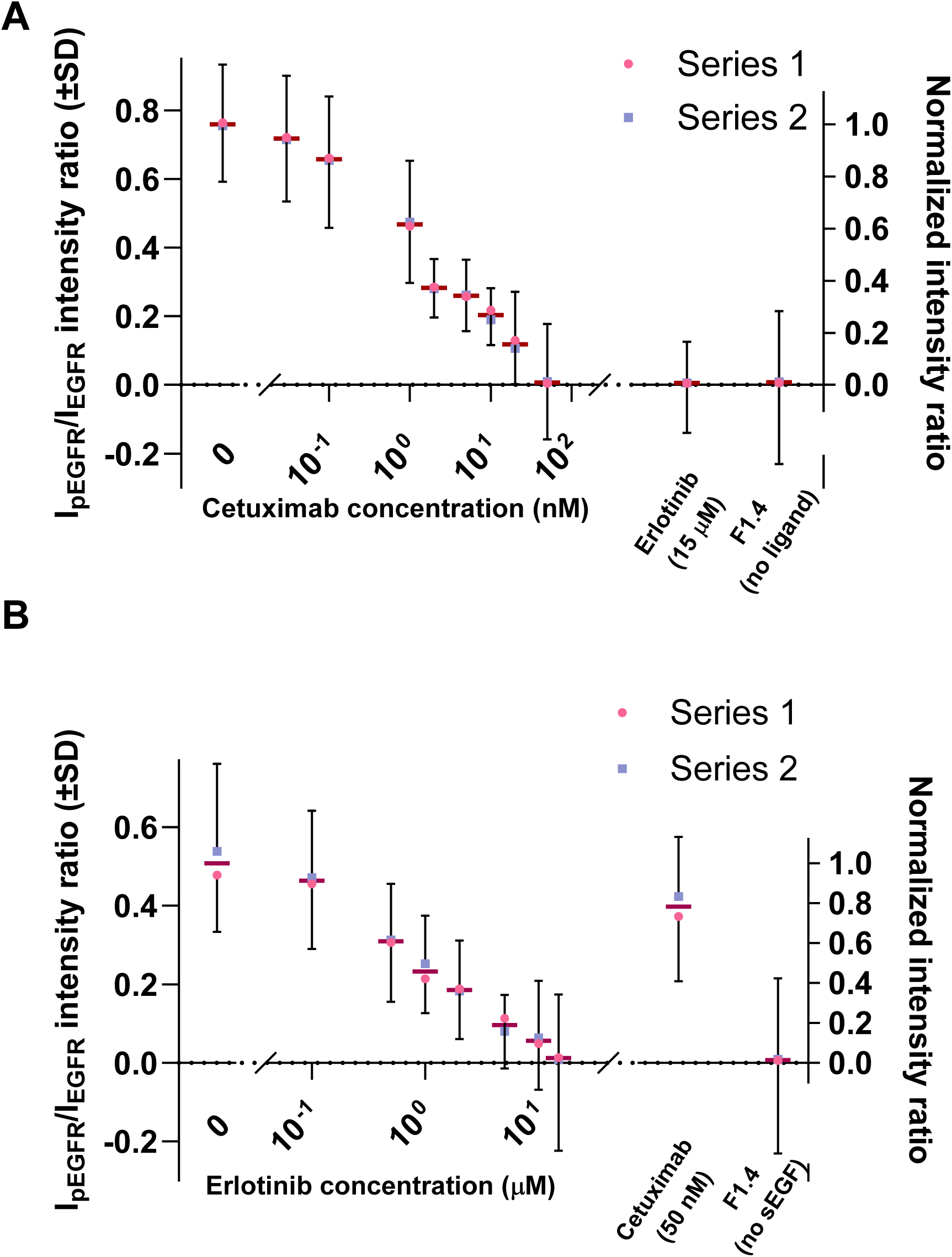
Effect of various concentrations of cetuximab extracellular and erlotinib intracellular inhibitors on EGFR phosphorylation in external EGF-treated or sEGF-producing F1.4 cells. (A) Fluorescence intensities of the Alexa 647-anti-pEGFR antibody (I_pEGFR_) and the EGFR-G (I_EGFR_) signals and their ratios were calculated for each cell. Averaged I_pEGFR_/I_EGFR_ intensity ratios (left y-axis) and their normalized values (right y-axis) for cells treated with extEGF (200 nM, 10 min) after pretreatment with different concentrations (0 to 50 nM, left x axis) of the EGFR inhibitor antibody cetuximab are shown on the left. 50 nM cetuximab completely inhibited receptor phosphorylation, similar to the intracellular kinase inhibitor erlotinib (15 µM, see on the right). No phosphorylation occurred in F1.4 cells in the absence of ligand (on the right). **(B)** Effect of erlotinib treatment (0-15 µM, on the left) on the I_pEGFR_/I_EGFR_ intensity ratios measured in F1.4 cells transfected with sEGF 24 h prior to inhibitor treatment. 15 µM erlotinib completely abolished EGFR phosphorylation. As a comparison, the extracellular inhibitor cetuximab (50 nM) reduced EGFR phosphorylation by ∼17%, it could not eliminate it completely (shown on the right). Cells not expressing sEGF showed no phosphorylation (on the right). Blue squares and pink circles represent averages from two independent experiments (Series 1 and 2), the upward and downward whiskers showing their SD-s. The burgundy lines in the middle represent the means of the corresponding averages in the two series. In each series 48-96 cells were evaluated.

The efficiency of phosphorylation and the efficacy of inhibitors was determined by calculating the cellular I_pEGFR_/I_EGFR_ intensity ratio, a quantity independent of the EGFR concentration (*Fig. 5)*. As reference, the intensity ratio measured in extEGF-treated F1.4 cells in the absence of inhibitors was used (leftmost point in Fig. 5A). All intensity ratios were normalized to this value; normalized values can be read from the right y axis. Increasing cetuximab concentrations gradually decreased phosphorylation efficiency. At the highest, 50 nM concentration, the normalized intensity ratio dropped to 0.009 indicating near-complete blocking of the effect of extEGF. Erlotinib, at the highest applied concentration (15 µM), was also used for this sample, reducing the intensity ratio to 0.01. F1.4 cells without any ligand or inhibitor treatment were used as a negative control resulting in a normalized intensity ratio of 0.009 (rightmost point in *Fig. 5A*). For sEGF transfected cells, erlotinib concentrations in the range of 0-15 µM were used. The highest absolute phosphorylation level induced by sEGF was I_pEGFR_/I_EGFR_=0.51 (its normalized value: 1.0), which decreased with increasing erlotinib concentrations. At 15 µM erlotinib, the normalized intensity ratio decreased to 0.02 showing that erlotinib was able to block EGFR activation in sEGF producing cells. On the contrary treating sEGF expressing cells with the highest applied concentration of cetuximab (50 nM), reduced the phosphorylation level only to 0.4 (*Fig. 5B*). This result implies that the cetuximab treatment is ineffective in inhibiting EGFR phosphorylation in cells producing the sEGF ligand.

## Discussion

EGFR plays a key role in both physiological and pathological signaling pathways within epithelial cells, facilitating a variety of biological responses through diverse signaling mechanisms. Soluble ligands of the EGFR generally activate the receptor via paracrine and autocrine pathways, while membrane-anchored ligands elicit juxtracrine signaling (Singh & Harris, 2005). Among the three primary signaling modes, autocrine signaling is particularly significant in the context of cancer development. In many tumor cells, which produce both EGFR and its ligand, autocrine loops are created, which results in persistent receptor activation at the plasma membrane. In tumor cells, the most common autocrine loops are formed by the co-expression of EGFR and its ligands EGF or TGFα. Such continual signaling drives uncontrolled proliferation and contributes to metastases and therapeutic resistance (Nickerson *et al*., 2013; Seth *et al*, 1999).

An additional mode of ligand-induced EGFR activation, termed intracellular autocrine or intracrine signaling, has been proposed. This signaling modality is a subgroup of autocrine signaling distinguished primarily by its subcellular localization, occurring not at the cell surface but within intracellular membrane compartments. Intracrine signaling of EGFR was first demonstrated by a research group, which showed that removal of the membrane-anchoring domain of preEGF caused both the ligand and receptor to relocate from the cell surface to intracellular vesicles. The growth of human mammary epithelial cells (HMEC) could be effectively inhibited by anti-EGFR antibodies when the membrane-anchoring domain was complete. In contrast, removing this domain caused the antibody treatment to become ineffective and enabled the colonies to continue growing. This implies that the autocrine signaling was converted to an intracrine form in HMEC cells making them resistant to extracellular antibody therapies (Wiley *et al*., 1998).

In line with these findings, we aimed to demonstrate the mechanistic steps of EGFR intracrine signaling with a focus on the Golgi apparatus. Using confocal microscopy, FLIM-FRET and FCS measurements, we traced the steps of intracrine signaling in a CHO cell model system, which lacks endogenous EGFR (Shi *et al*., 2000). The initial steps in this process involve ligand binding, then dimerization of EGFR chains and subsequent phosphorylation of the tyrosine residues (Singh & Harris, 2005). We summarized our results in a putative model of intracrine signaling in the Golgi in *Fig. 6*. A previous study (Wiley *et al*., 1998) observed the colocalization of sEGF and EGFR in intracellular vesicles at the resolution of light microscopy (∼200 nm); however, the molecular interaction was not directly detected. We assessed the binding of both the precursor and soluble forms of EGF by FLIM-FRET, which can report on molecular vicinity in the 2-10 nm range (Orthaus, 2009). FLIM-FRET measurements revealed that sEGF is found in the proximity to EGFR already in the Golgi, visualized by the B-Giantin protein. Conversely, preEGF and EGFR showed no interaction within the Golgi or in other compartments of the cell (*Fig 1B.*). The lack of interaction is probably due to steric hindrance between the N-terminal extension of preEGF and EGFR ligand pocket (Dong *et al*, 2005). In contrast, the heparin binding EGF and transforming growth factor-α have biological activity also in their membrane-anchored precursor form and can activate receptors in a neighboring cell’s membrane in a juxtracrine manner (Bush *et al*, 1998; Dong *et al*., 2005; Iwamoto & Mekada, 2000). The very low FRET efficiencies detected between preEGF and EGFR were equal with those in the negative controls indicating that these E values can result from random spatial colocalization rather than specific receptor-ligand interactions (King *et al*, 2014). The distribution of EGFR also changed in sEGF expressing samples: it was mainly located within intracellular vesicles, whereas it was found mainly in the cell membrane in preEGF producing cells (*Fig. 1A).* These results are aligned with similar observations described in (Wiley *et al*., 1998). To confirm sEGF ligand binding with an independent method, FCS was utilized, where the mobility of G-sEGF was measured in the presence or absence of EGFR, along with freely diffusing and membrane-bound controls, eGFP and G-EGFR, respectively. G-sEGF mobility parameters measured in the absence of EGFR were similar to those of the freely diffusing eGFP. However, when co-expressed with EGFR, the mobility of G-sEGF was significantly slower approximating that of G-EGFR (*Fig. 2B*). These results also suggest that at least a fraction of the sEGF ligand binds to the receptor.

**Fig. 6.**
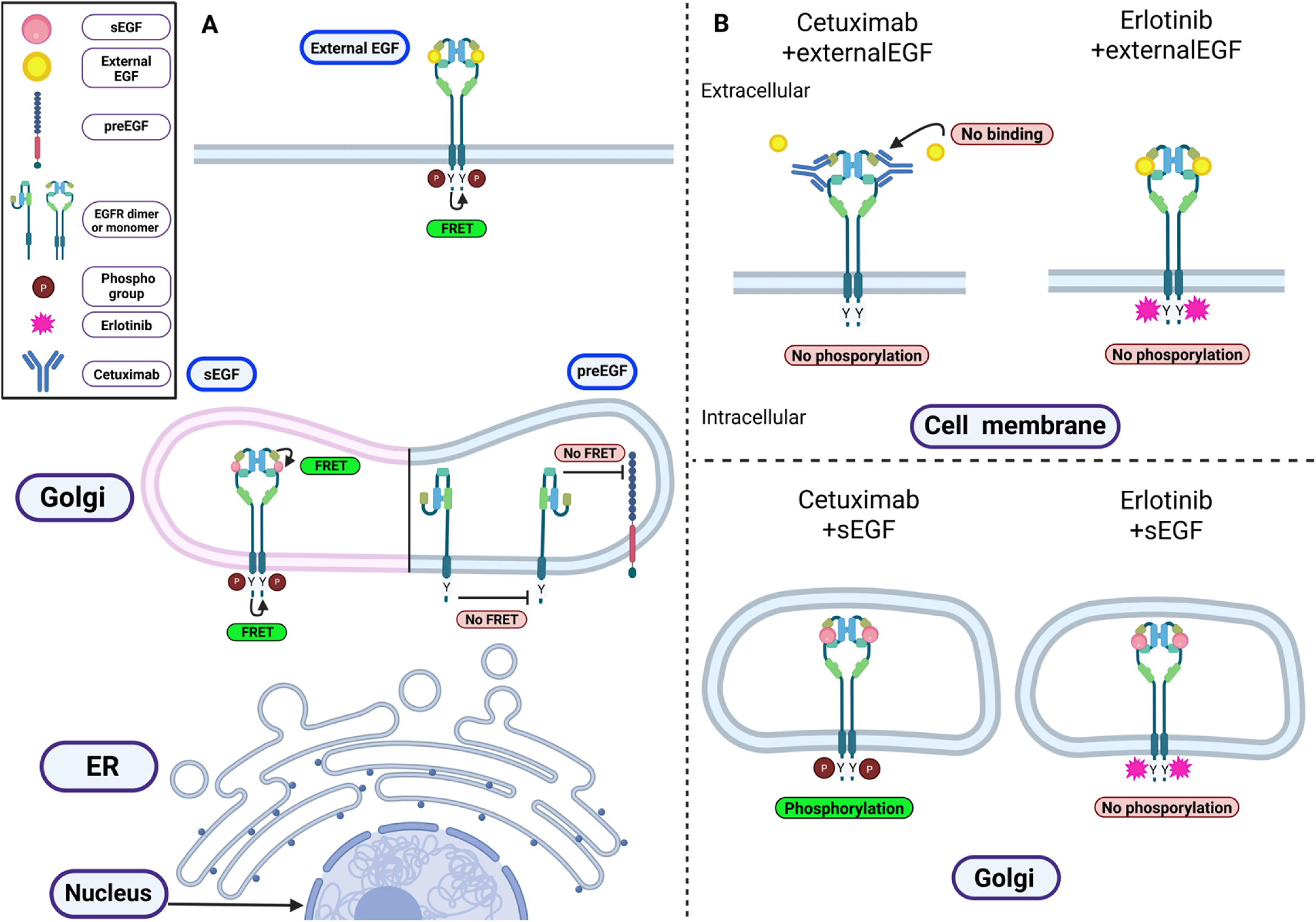
(A) Schematic representation of the intracrine signaling process of EGFR. External EGF binds to EGFR at the plasma membrane inducing its dimerization and activation. sEGF binds to the EGFR in the Golgi, initiates its dimerization and phosphorylation in the Golgi, while preEGF cannot bind. **(B)** Schematic representation of the effect of extracellular cetuximab and intracellular erlotinib inhibitors on EGFR phosphorylation. In F1.4 cells treated with external EGF ligand and cetuximab or erlotinib, no EGFR phosphorylation was observed. F1.4 cells transfected with sEGF, the cetuximab could not eliminate the phosphorylation as not entering into the Golgi. In contrast, the intracellularly acting erlotinib eliminates EGFR phosphorylation in these cells (Image created with Biorender.com).

To test whether the receptor chains can also preassemble already in the Golgi, we also used FLIM-FRET. The results indicated at least a partial association between donor-and acceptor-tagged EGFR chains in the Golgi of cells producing sEGF (*Fig. 3*); for preEGF, no significant association was detected. The FRET efficiency between receptors was also low in the cell membrane of preEGF producing or ligand free control cells. It has been a long-standing question whether EGFR in the plasma membrane was monomeric or dimeric/oligomeric prior to activation. EGFR was found in a mixture of these states at different conditions and in different cell lines (Balasubramanian *et al*, 2022; Hajdu *et al*., 2020; Sankaran *et al*, 2021; Yavas *et al*, 2016). In our system, the receptor was predominantly monomeric in the absence of ligand whereas external EGF treatment enhanced the degree of dimerization/oligomerization in the membrane to a level similar to that measured in the Golgi of cells expressing sEGF (*Fig. 3B*).

Immunofluorescence labelling of pEGFR was carried out on F1.4-B-Giantin cells co-expressing EGFR and sEGF or preEGF to detect receptor activation within the Golgi. A high level of phosphorylation was observed in intracellular vesicles containing both sEGF and EGFR. In contrast, cells expressing preEGF exhibited a low, homogeneously distributed background phosphorylation signal similar to control cells and in cell expressing only EGFR. Pearson’s correlation coefficients indicated a strong colocalization between EGFR and sEGF in the Golgi, while for preEGF only a weak correlation was detected with EGFR.

Finally, the effects of the extracellular inhibitor cetuximab and the intracellular inhibitor erlotinib were tested on EGFR phosphorylation. Earlier it was reported that the proliferation of sEGF producing cells could not be inhibited by extracellular blocking antibodies (Wiley *et al*., 1998). In our experiments, both cetuximab and erlotinib reduced phosphorylation to almost zero in a concentration-dependent manner in F1.4 cells treated with extEGF (but not expressing ligands). On the other hand, in sEGF producing cells, 15 µM erlotinib (the highest concentration applied for F1.4) blocked phosphorylation completely while cetuximab, even at its highest concentration (50 nM), diminished EGFR phosphorylation by only ∼20%. This partial efficiency may be explained by inhibition of those EGFR’s that had not bound sEGF and had been trafficked to the cell membrane. These findings altogether imply the presence of intracrine EGFR signaling in sEGF expressing cells, which cannot be inhibited by extracellularly blocking antibodies. We summarized our results graphically in *Fig. 6B*.

Previously, we investigated the mechanism and steps of the intracrine signaling of the interleukin-2 receptor (IL-2R) (Volko *et al*., 2019). In a HeLa cell model system, the subunits of the heterotrimeric IL-2R were partially preassembled in the ER and the Golgi. In adult T-cell leukemia (ATL) cells producing both IL-2R and its ligand, receptor and JAK1/3 phosphorylation already occurred in the Golgi apparatus. A combination of antibodies targeting IL-2 and membrane-bound IL-2Rα and IL-2/15Rβ was ineffective to inhibit proliferation of cells expressing their own IL-2 while IL-2 non-expressing cells were efficiently blocked. Importantly, IL-2 is synthesized as a soluble peptide whereas preEGF is produced as an inactive membrane-bound precursor, which, in its soluble form, also induces intracrine signaling in the same manner. For EGFR, receptor activation is controlled at several levels from receptor and ligand expression to the presence and activity of metalloproteinases involved in ligand shedding (Ohtsu *et al*., 2006; Sahin *et al*, 2004). Notably, the generation of sEGF in the Golgi apparatus requires the shedding of the pre-EGF ligand by ADAM enzymes; however, such a process has not yet been observed and needs further investigation.

Our finding that cetuximab was ineffective in cells expressing sEGF while erlotinib effectively blocked EGFR activation brings up the possibility that intracrine signaling may contribute to cetuximab resistance. The enzyme ADAM17 is often overexpressed in many tumors (e.g., breast, non-small cell lung, colorectal, and gastric cancers), where it promotes continuous tumor cell proliferation by activating EGFR through the shedding of its membrane-bound precursor ligands, contributing to poor cancer prognosis and resistance (Arribas & Esselens, 2009; Chen *et al*, 2024; Li *et al*, 2019; Ni *et al*, 2013; Saha *et al*, 2022). Acquired resistance to extracellular cetuximab/panitumumab therapies in metastatic tumor cells overexpressing EGFR often arises through diverse, complex pathways that enable these cells to evade inhibition. Modified versions of mAbs, such as cetuximab conjugated to ribonuclease A (RNase A), which combine extracellular and intracellular blocking, had promising results in signal transducing KRAS protein mutated colorectal cancer cells, where RNase A caused RNA degradation upon its release in the cytosol and triggered apoptosis (Jafary *et al*., 2025). Other studies have examined the use of tyrosine kinase inhibitors (TKIs) either alone or in combination with monoclonal antibodies, as potential treatment strategies for metastatic cancers. Erlotinib showed a promising effect in cetuximab resistant NSCLC cells (Brand *et al*, 2011a), while afatinib was effective in colorectal cancer (De Pauw *et al*, 2019). In human colorectal cancer cells, crizotinib, a tyrosine kinase inhibitor simultaneously targeting receptors like EGFR, MET (HGFR, Hepatocyte growth factor receptor), and RON (MST1R, macrophage-stimulating protein receptor), effectively overcame cetuximab resistance in both *in vitro* 3D cell culture models and *in vivo* mouse xenografts (Graves-Deal *et al*, 2019).

In conclusion, our paper provides compelling evidence that in cells with high endogenous sEGF production, EGFR associates with sEGF in the Golgi leading to its dimerization and activation before reaching the plasma membrane. Although this intracrine signaling route may be less significant under normal conditions, it may become highly relevant in tumor cells with EGFR overexpression and abnormal ligand processing. This intracrine signaling pathway may sustain the continual activation of cells rendering extracellular blockade by monoclonal antibodies to become ineffective. The reduced expression of EGFR may also decrease ADCC. Therefore, effective therapeutic strategies may require the use of intracellular kinase inhibitors alone or in combination with monoclonal antibodies or novel agents targeting intracellular pathways to overcome acquired resistance and improve clinical outcomes.

## Materials and Methods

### Cell culture

Chinese Hamster Ovary (CHO), the CHO-derived subline F1.4 and Human embryonic kidney (HEK293T) cell lines were cultured in Dulbecco’s Modified Eagle Medium (Invitrogen Life Technologies, Carlsbad, CA) supplemented with 10 % (v/v) fetal-bovine serum (FBS, Sigma-Aldrich), 100 U/ml penicillin and 100 mg/ml streptomycin. The F1.4 cell line stably expresses 1-2 million copies of EGFR C-terminally tagged with eGFP (Brock *et al*, 1999). The F1.4-B-Giantin cell line is a version of F1.4 transduced with TagBFP-Giantin as well. For virally transduced cell lines DMEM was additionally supplemented with puromycin (1.2 mg/ml) or geneticin (0.6 mg/ml or 1.2 mg/ml) for single-transduced or with both for double-transduced cells. All cells were maintained in a 5% CO_2_-humidified atmosphere at 37°C.

### Plasmid construction

The plasmid encoding Giantin tagged with BFP (referred to as BFP from hereon), a Golgi-resident protein, was previously generated by our research group as described in (Volko *et al*., 2019). As a negative control for FLIM-FRET eGFP-C3 (Clontech Laboratories, Mountain View, CA) and mScarlet3-C3 vectors were used. The mScarlet3 fluorescent protein was amplified from the pC1-mScarlet3-iii vector, a gift of Prof. T. Gadella (University of Amsterdam, the Netherlands), and ligated into the eGFP-C3 vector using AgeI/BsrGI restriction enzymes. The mScarlet3-IL-2Rα construct was created by replacing the mCherry fluorescent protein in mCherry-IL-2Rα with mScarlet3. The eGFP-mScarlet3 fusion protein containing a four-amino-acid linker (YSDL) was generated by excising mCherry from the eGFP-mCherry plasmid with BgII and SalI enzymes and replacing it with mScarlet3. The plasmid encoding eGFP-EGFR was constructed in two steps: first, the EGFR signal peptide was subcloned into the eGFP-C3 vector using the NheI/AgeI enzymes; second, the sequence downstream of eGFP was replaced with the EGFR coding sequence amplified from the hEGFR-pcDNA3 construct (a kind gift from Donna Arndt-Jovin, Max Planck Institute for Multidisciplinary Sciences, Göttingen, Germany). For the mScarlet3-EGFR vector, eGFP was excised using XhoI/HindIII and replaced with mScarlet3. To measure receptor dimerization, a plasmid encoding C-terminally mScarlet3-tagged EGFR was created in the mScarlet3-N1 transient vector via NheI/AgeI restriction sites (the mScarlet3-N1 vector was originally derived from the eGFP-N1 vector, Clontech Laboratories). Subsequently, mScarlet3 was excised using AgeI/BsrGI and replaced with eGFP to create the EGFR-eGFP construct.

The cDNA encoding preEGF (OriGene, SC127840) was used to generate the N-terminally mScarlet3-tagged sEGF/preEGF ligands in two steps. First, the signal peptide sequence was inserted with NheI/AgeI restriction sites to the N-terminus of the mScarlet-3-C3 vector. In the second step, the remaining coding sequence of the sEGF/preEGF ligand was cloned to the C-terminus using XhoI and SalI. This mScarlet3-sEGF construct was used to create the eGFP-sEGF and BFP-sEGF plasmids, where the mScarlet3 tag was removed and replaced with eGFP or BFP, respectively. Similarly, the mScarlet3-preEGF ligand was processed to produce the BFP-labeled version. To clone the preEGF-mScarlet3, the full DNA sequence of the signal and the ligand was amplified by PCR and ligated into the mScarlet3-N1 vector with the XhoI/SalI sites. The backbone for the HaloTag-Giantin construct was generated from the eGFP-C3 plasmid by removing eGFP with AgeI and BglII and replacing it with the HaloTag. In a following step, the Giantin was subcloned with the BamHI/KpnI sites.

To establish cell lines stably expressing BFP-Giantin, mScarlet3-preEGF, preEGF-mScarlet3, BFP-sEGF and BFP-preEGF, the second-generation viral transfer vector, pBMN-Z-IN (Addgene plasmid #1735, a gift from Garry Nolan) was used. The pBMN-Z-IN-puro-IRES-IL2Rβ vector with puromycin resistance was created earlier by replacing the geneticin resistance coding gene with puro-IRES-IL2Rβ in pBMN-Z-IN. The BFP-Giantin and Halotag-Giantin inserts were subcloned into pBMN-Z-IN-puro-IRES-IL2Rβ vector with MauBI and SalI restriction enzymes by excising the IL2Rβ. The sequence of the plasmids was confirmed by Sanger sequencing (UD-GenoMed, Debrecen, Hungary).

### Viral transduction and stable cell lines

The HEK293T packaging cell line was used to produce viral particles for generating stable cell lines expressing genes of interest. Both CHO and F1.4 cell lines were stably transfected using the calcium phosphate transfection method with VSVG and PAX2 helper plasmids. To visualize the Golgi, the pBMN-Z-IN-BFP-Giantin (geneticin resistance) or pBMN-Z-IN-puro-IRES-BFP-Giantin (puromycin resistance) retroviral vectors were transfected. Established F1.4-BFP-Giantin and CHO-BFP-Giantin lines with puromycin resistance were subsequently transduced with pBMN-Z-IN-mScarlet3-preEGF and pBMN-Z-IN-preEGF-mScarlet3 constructs. For EGFR dimerization experiments, CHO cells were first transduced with pBMN-Z-IN-puro-IRES-HaloTag-Giantin and later stably transfected with pBMN-Z-IN vectors encoding BFP-sEGF or BFP-preEGF.

Retroviral vectors containing the appropriate insert(s), helper plasmids, 2× HEPES-buffered saline, and CaCl₂ were combined and added to HEK293T cells along with 12 µl of 25 mM chloroquine. Forty-eight hours later, the supernatant containing the assembled virus particles was harvested, filtered through a 0.45 µm syringe filter, after which the cells of interest were transfected with it. The efficiency of viral transduction was increased by the addition of 10 µg/ml polybrene (Sigma-Aldrich) and checked after 72 hours using a confocal microscope. Cell lines were sorted using a CytoFLEX SRT Cell Sorter (Beckman-Coulter, Brea, USA) to separate the transduced cells from the non-transduced ones.

### Transient transfection

In the experiments 6000-9000 cells/well were plated in chambered coverslips (ibidi) containing 300 µl 10% FBS DMEM and the appropriate antibiotics for selection. The transfection of the seeded cells with FuGene HD or FuGene 4K (both from Promega, Madison, WI) reagents were carried out 24 hours later. In the reactions 0.4 µl FuGene HD or 0.6 µl FuGene 4K was mixed with 100-120 ng plasmid. We optimized the amount of DNA for co-transfection to achieve equal expression levels. Relative expression levels were assessed by comparing the brightness ratio of the fluorescent proteins in co-transfected cells to that arising from the appropriate fusion protein (such as eGFP-mScarlet3). Cells were measured 24-48 hours after transfection and, prior to measurements, the medium was replaced with Leibovitz’s L-15 imaging medium (Thermo Fisher Scientific).

### Immunofluorescence labeling

Cells grown in an 8-well chamber (ibidi) were starved for 4 hours at 37 °C, where the medium was replaced with serum-free 10 mM glucose containing HEPES buffer. After starvation, ibidi chambers were placed on an ice-cold parafilm-coated metal surface and washed twice with ice-cold HEPES buffer. Then cells were fixed with 2% formaldehyde-HEPES for 60 minutes at 4 °C. After fixation, 300 µl 100% methanol was added for 5 minutes. This was followed by a washing step with HEPES buffer twice. For permeabilization 300 µl 0.1% TritonX-100/1%BSA solution was used for 5 minutes. The blocking step included one hour incubation with 1% BSA-HEPES solution at room temperature. The rabbit recombinant mAb against the phosphorylated Y1173 residue of EGFR conjugated with AlexaFluor-647 (A647, ab207346, Abcam) was diluted in 0.05% Triton-X100 1% BSA-HEPES solution at a 1:200 ratio. Cells were incubated with the antibody overnight at 4 °C. The following day, the cells were washed twice in HEPES buffer solution, after which they were fixed with 200 µl of 2% formaldehyde for 30 minutes at 4 °C prior to confocal imaging.

### Confocal fluorescence microscopy

Confocal fluorescence microscopy was performed to capture images used for calculating Pearson’s correlation coefficients to describe the colocalization of pre/sEGF and EGFR in the Golgi, and to assess the effect of extracellular and intracellular inhibitors on the phosphorylation level of EGFR. The confocal images were taken by an A1 confocal microscope (Nikon, Tokyo, Japan) equipped with a Plan-Apochromat 60x water immersion objective, NA 1.27. The zoom factor was set to 2 for quantifying pEGFR levels and 3.5 for images used in Pearson’s coefficient calculations. The pinhole size was adjusted to 1.2 Airy units. Fluorophores BFP, eGFP, mScarlet3, and A647 were excited using 405 nm, 485 nm, 561 nm, and 647 nm lasers and detected through 430/475, 500/550, 570/620 and 663/738 corresponding bandpass filters, respectively.

Distinct false colors were assigned to each channel: blue for BFP-Giantin, green for eGFP-EGFR, red for mScarlet3-tagged pre/sEGF, and magenta for anti-pEGFR-mAb-A647. To visualize overlap in composite images, the mAb channel false color was changed from magenta to green, resulting in white color representing triple colocalization between the Golgi, the ligand, and pEGFR.

### Evaluation of colocalization by Pearson’s correlation coefficient

Pearson’s correlation coefficient (PCC) was used to quantify colocalization of receptor chains within the Golgi apparatus. The calculations were performed using a customized MATLAB (Mathworks Inc, Natick, MA) program using DIPimage function (Delft University of Technology, Delft, The Netherlands). Confocal images of eGFP-EGFR, mScarlet3-s/pre-EGF, and anti-pEGFR-mAb-A647 were acquired alongside the BFP-Giantin Golgi marker. The region of interest (ROI) was defined by manual thresholding in the Golgi channel, and this mask was applied to all other channels to restrict colocalization analysis to Golgi-localized pixels.

The PCC was determined for the following pairs: EGFR/preEGF or EGFR/sEGF ligand. Detectable phosphorylation occurred only in samples expressing sEGF; therefore, additional PCC analyses were only performed for samples expressing sEGF but not for preEGF, namely, the Golgi/anti-pEGFR mAb, sEGF/anti-pEGFR mAb and EGFR/anti-pEGFR mAb pairs.

The principle of the calculation of the PCC has been described previously (Costes *et al*, 2004). As a negative control, the PCC was calculated for two images, in one of which pixels were randomly shuffled while leaving the pixels in the other image unscrambled. This process allows to distinguish true colocalization from random overlap. The process was repeated 100 times to generate a distribution of correlation coefficients representing the absence of correlation. If the calculated correlation coefficient value for the original image pair fell outside the 95% confidence interval of the distribution, colocalization was considered statistically significant. 51-74 cells were evaluated from each sample.

### Ligand and inhibitor treatments

In dimerization measurements the cells were treated for 2 hours with 10 µM of erlotinib (Selleckchem, Houston, TX, United States) kinase inhibitor to prevent internalization of EGFR (Hajdu *et al*., 2020). Next, we used a 200 nM of the external EGF (referred to as extEGF in the results, E9644-5MG, Sigma-Aldrich) ligand for five minutes at 37 °C in a 5% CO_2_ incubator to activate the EGFR. This was followed by the fixation of cells with 2% formaldehyde for 30 minutes at 4 ℃ prior to FLIM-FRET.

SiR HaloTag ligand (Lukinavicius *et al*., 2013) is a cell permeable compound, which was used to label the HaloTag-Giantin construct to visualize the Golgi. The stock solution with 20 mM concentration was diluted in two steps with DMSO to 600 µM, from which 1.5 µl was added to the cells in 300 µl DMEM (3 µM final concentration). Cells were incubated at 37 °C, 5% CO_2_ with the ligand for 1 hour and then washed with warm medium for 1 hour at the same conditions.

To study blocking of EGFR phosphorylation, various concentrations of cetuximab (Erbitux, Merck Serono, Darmstadt, Germany; 0, 0.03, 0.1, 1, 2, 5, 10, 20 and 50 nM) and erlotinib (0, 0.1, 0.5, 1, 2, 5, 10 and 15 µM) were used for 2 hours in F1.4 cells. Next, the cells were treated with 200 nM of extEGF ligand for 10 minutes at 37 ℃ before immunostaining.

### Evaluation of the inhibitory effect of cetuximab and erlotinib on EGFR phosphorylation

For testing the effect of inhibitors, we used F1.4 cells transfected with mScarlet3-sEGF 24 hours before inhibitor treatments. We also used non-transfected F1.4 cells. Both samples were treated the next day with extracellular cetuximab and intracellular erlotinib inhibitors as described above. Following inhibitor treatment, the non-transfected sample was stimulated with 200 nM of extEGF for 10 minutes. Both samples were fixed and labelled with the pEGFR-mAb-A647.

First, confocal images were acquired in EGFR-eGFP, mScarlet3-sEGF, and anti-pEGFR-mAb-A647 channels. ROIs were defined for the whole cell by manually outlining individual cells on the transmission image using ImageJ (Schneider *et al*, 2012), and these ROIs were then applied to the fluorescence channels corresponding to sEGF and EGFR. Background autofluorescence for each channel was determined from “empty” CHO cells and subtracted from the measured intensities. The pEGFR/EGFR intensity ratios were calculated from background-corrected intensities on a cell-by-cell basis, and the mean ratios were plotted at each inhibitor concentration. Normalizing pEGFR fluorescence intensity with the EGFR intensity makes this parameter an expression level independent measure of phosphorylation efficiency. In every sample 48-96 cells were analyzed.

### Fluorescence correlation spectroscopy (FCS) measurement

Fluorescence correlation spectroscopy (FCS) serves as a great tool to study molecular interactions and dynamics in various environments. Hence, we can study the receptor-ligand binding in living cells. FCS measurement is based on the fluorescence signal fluctuations caused by fluorescently tagged molecules diffusing in and out of the sub-femtoliter confocal detection volume (R. Rigler, 1993). From these fluctuations the autocorrelation function (ACF) is calculated, which is fitted with different models to determine the fractions of molecular subpopulations and their diffusion times or diffusion coefficients (Brock *et al*., 1999; Vamosi *et al*, 2009).

The mobility of eGFP alone and eGFP-EGFR/eGFP-sEGF was measured with a Zeiss LSM880 confocal laser scanning microscope (Carl Zeiss, Jena) using a 40x water immersion objective (NA 1.2). First, we recorded confocal images of cells, where the BFP-Giantin was excited with a 405-nm diode laser and its blue fluorescence signal was detected between 415 and 490 nm; eGFP was excited by the 488-nm line of an Argon-Ion laser and detected between 500 and 534 nm; mScarlet3-EGFR was excited by a HeNe laser at 543 nm and the signal detected between 578 and 696 nm. To measure autocorrelation curves of the eGFP-tagged constructs the 488-nm line was used. For each cell, 10 runs 8 seconds each were recorded at selected points in the Golgi. All measurements were carried out at room temperature (23 °C). To select cells with an acceptor-to-donor ratio around 1, we used the eGFP-mScarlet3 fusion protein as a control.

The FCS data were analyzed with the Quickfit 3.0 software (https://biii.eu/quickfit-3). All the runs in the ACF curves were inspected, and those with artifacts (e.g., due to cell movement or large photobleaching) were excluded from the calculations. The ACFs from the remaining runs were averaged and then fitted with a normal diffusion model (with particles diffusing in 3D or 2D) to extract the fractions and diffusion coefficients of the individual subpopulations. Fitting was performed with a simulated annealing algorithm with box constraints was, using a weighting model based on the standard deviations of the runs.

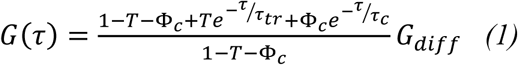

where

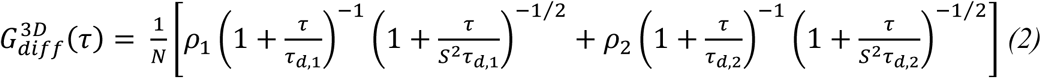

and

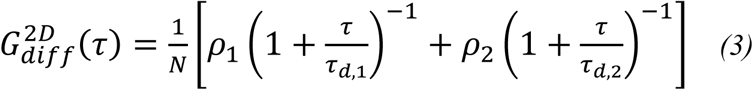

where N indicates the average number of fluorescent molecules present within the detection volume, τ_TR_ is the triplet correlation time (fixed to 3 µs), T shows the equilibrium molar fraction of fluorophores in the triplet state and τ is the lag time (Widengren & Rigler, 1998). eGFP undergoes two distinct protonation processes: an intramolecular proton transfer and a pH-dependent extracellular protonation (Haupts *et al*, 1998). Because the difference between the characteristic time constants of the two protonation processes at pH 7.4 is less than one order of magnitude, they were modeled together as a single term, characterized by the molecular fraction Θ_c_ and a fixed correlation time τ_c_ of 320 µs (Vamosi *et al*., 2019). The S value, which represents the ratio of the axial to longitudinal diameters of the ellipsoidal detection volume, was obtained at the beginning of the measurement process through the fitting the ACF of the Alexa Fluor 488 solution (100 nM in 10 mM Tris-EDTA, pH 7.4).

To calculate the diffusion coefficients D_i_ in the samples the following formula was used:

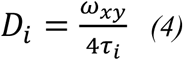

where ω_xy_ is the lateral radius of the detection volume. ω_xy_ was determined from the diffusion time of the Alexa Fluor 488 dye as:

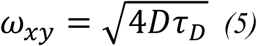

where τ_D_ is the measured diffusion time of the A488 and D is the diffusion coefficient of the dye, 414 µm^2^/s at 25 °C (Petrov *et al*, 2006). In both the 2D or 3D models, two populations were considered: a fast component with fraction ρ₁ and diffusion time τ₁, and a slow component with fraction ρ₂ and diffusion time τ₂, where ρ₂ = 1 − ρ₁. We measured 36-53 cells from each sample.

### FLIM-FRET measurement

Förster Resonance Energy Transfer (FRET) measured by fluorescence lifetime imaging microscopy (FLIM) is a powerful tool to investigate molecular interactions such as homodimerization/oligomerization of receptor chains or receptor-ligand binding. FRET is the non-radiative transfer of excitation energy between an excited donor and an acceptor fluorophore that are within each other’s molecular vicinity (2-10 nm) through dipole-dipole coupling. The efficiency of this process (FRET efficiency, E) depends inversely on the sixth power of the distance between the donor and acceptor (Förster, 1948). In our experiments we used eGFP as a donor and mScarlet3 as an acceptor. The lifetime of the donor decreases in the presence of the acceptor fluorophore, which forms the basis of FLIM-based detection of E.

Measurements were performed using a Nikon A1 confocal microscope equipped with a time-correlated single-photon counting (TCSPC) upgrade (Picoquant, Berlin, Germany). Before all measurements, preview images were taken with the 405 nm, 485 nm, 561 nm and 647 nm CW lasers and detected through the 430/475, 500/550, 570/620 and 663/738 bandpass filters for visualizing the fluorescent proteins and later calculating the acceptor-to-donor expression ratio (N_A_/N_D_). Unique pseudocolors were selected for each channel. In pre/sEGF-EGFR binding experiments, blue indicated the BFP-Giantin Golgi marker, green the eGFP-tagged EGFR or sEGF, and red the mScarlet3-tagged pre/sEGF ligand or EGFR. For dimerization studies, blue represented BFP-labeled pre/sEGF, green/red the eGFP/mScarlet3-tagged EGFR chains, and magenta denoted the SIR-HaloTag-labeled Giantin Golgi marker. We created composite images of the Golgi and ligand channels alongside the EGFR channel to identify areas of overlap, which appeared in white color. The magenta color of SIR-tagged Golgi was changed to red in the overlap image, and it was used in combination with the EGFR-eGFP and BFP-s/preEGF to result in white color for triple overlap.

For FLIM measurements, eGFP was excited with a 485-nm picosecond pulsed laser with 30-35 MHz repetition rate and emission was detected with a PMA hybrid 40 photon-counting photomultiplier (PicoQuant) through a 520/30 nm emission filter. BFP-Giantin images were recorded using a 400/40 nm, while SIR-Halo-tag-Giantin via a 670/90 nm filter. Data were collected for 10 s for the Golgi marker (BFP or SIR-HaloTag-Giantin) and 60 s for the eGFP donor. The preview image of the acceptor (mScarlet3) was recorded prior to the FLIM measurement with the CW laser of the A1 confocal microscope. All FLIM-FRET measurements were performed at room temperature (23 °C).

### Evaluation of FLIM-FRET

To analyze the fluorescence decay curves, the SymphoTime 64 software (PicoQuant) was used. Prior to determining fluorescence lifetimes in the Golgi or the plasma membrane, images were intensity thresholded to exclude background pixels and then the Golgi apparatus or the membrane was selected with a freehand-drawn ROI. For the eGFP-mScarlet3 fusion (positive control) and the co-expressed eGFP and mScarlet3 (negative control), the ROI was drawn around the entire cell, excluding the nucleoli. The decay curves for the selected ROIs were fitted to a multiexponential re-convolution model with two lifetime components:

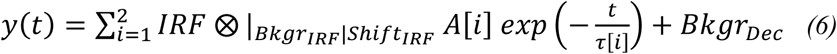

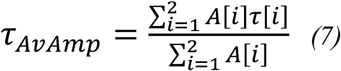

where and τ[i] and A[i] are the exponential decay time and the amplitude of the i^th^ component, Bkgr_Dec_ is the background correction, Bkgr_IRF_ is the background correction for the instrument response function (IRF), Shift_IRF_ is the correction for temporal IRF displacement, and τ_Av,amp_ amplitude-weighted average lifetime.

The FRET efficiency was calculated in the Golgi area/plasma membrane with the following formula:

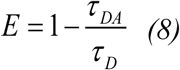

where τ_DA_ is the amplitude-weighted average lifetime of the donor in the donor-acceptor-containing samples, and τ_D_ is that in the absence of the acceptor molecule (Orthaus, 2009).

For selected cells, in the defined ROI, the pixel-by-pixel distribution of amplitude-weighted average lifetime map of the donor, the FRET efficiency map and the FRET efficiency histogram were calculated. The eGFP-mScarlet3 fusion protein was measured as a positive control, which expresses the donor and the acceptor at a 1:1 ratio. The FRET-corrected fluorescence intensity ratio for the positive control (Q) was calculated using the following formula:

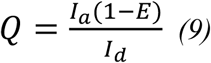

Where I_A_ is the intensity of the acceptor fluorophore, I_D_ is the intensity of the donor, E is the FRET efficiency. The I_a_ and I_d_ intensity values were determined from the representative confocal images by applying ROI for the whole cell, Golgi or membrane in ImageJ software (Schneider *et al*., 2012). For other samples, the acceptor-to-donor molecular ratio, N_a_/N_d_, was quantified as (Vamosi *et al*, 2008).

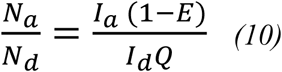

## Statistical analysis

Statistical analysis of the different measurements was carried out with the GraphPad Software (version 10.0.0). Mean values were compared with unpaired t-tests. Data are presented as scatter dot plots showing mean% ± standard deviation (SD)% for FLIM-FRET measurements. For FCS measurements, results are displayed as box plots, where the middle line represents the median, the top and bottom edges of the box correspond to the 75^th^ and 25^th^ percentiles, respectively, and the whiskers indicate the 90^th^ and 10^th^ percentiles.

## Author contributions

Noémi Bilakovics: Conceptualization, Formal analysis, Investigation, Methodology, Validation, Visualization, Writing - original draft, Writing - review & editing. Julianna Volkó: Investigation, Methodology, Formal analysis. Chinzorig Jeppesen: Investigation, Formal analysis. Gergely Fekete: Investigation, Formal analysis. István Csomós: Investigation, Formal analysis. Andrea Bodnár: Editing original draft. Krisztina Németh: Resources. Attila Kormos: Resources. Katalin Tóth: Editing original draft, Conceptualization. György Vereb: Conceptualization, Editing original draft. Péter Nagy: Conceptualization, Editing original draft, Formal analysis. György Vámosi: Conceptualization, Funding acqusition, Project administration, Supervision, Validation, Methodology, Writing-original draft& editing, Writing - review & editing.

## Acknowledgements

We thank Edina Nagy for her excellent technical assistance. Microscopy measurements were carried out in the Sándor Damjanovich Cell Analysis Core Facility of the University of Debrecen (Cellular BioImaging Hungary, a Euro-BioImaging Node). This work was supported by NKFIH-OTKA grants K129371, ANN 135107, K146028, ANN133421, K138075, HUN-REN Hungarian Research Network, Lendület “Momentum” Program of the Hungarian Academy of Sciences (LP-2024-11) and by 2024-1.2.2-ERA_NET-2024-00009 from the National Research, Development and Innovation Office, Hungary. Supported by the University of Debrecen Program for Scientific Publication.

